# Ribosome Decision Graphs for the Representation of Eukaryotic RNA Translation Complexity

**DOI:** 10.1101/2023.11.10.566564

**Authors:** Jack A. S. Tierney, Michał Świrski, Håkon Tjeldnes, Jonathan M. Mudge, Joanna Kufel, Nicola Whiffin, Eivind Valen, Pavel V. Baranov

## Abstract

The application of ribosome profiling has revealed an unexpected abundance of translation in addition to that responsible for the synthesis of previously annotated protein-coding regions. Multiple short sequences have been found to be translated within single RNA molecules, both within annotated protein-coding and non-coding regions. The biological significance of this translation is a matter of intensive investigation. However, current schematic or annotation-based representations of mRNA translation generally do not account for the apparent multitude of translated regions within the same molecules. They also do not take into account the stochasticity of the process that allows alternative translations of the same RNA molecules by different ribosomes. There is a need for formal representations of mRNA complexity that would enable the analysis of quantitative information on translation and more accurate models for predicting the phenotypic effects of genetic variants affecting translation. To address this, we developed a conceptually novel abstraction that we term Ribosome Decision Graphs (RDGs). RDGs represent translation as multiple ribosome paths through untranslated and translated mRNA segments. We termed the later ‘translons’. Non-deterministic events, such as initiation, re-initiation, selenocysteine insertion or ribosomal frameshifting are then represented as branching points. This representation allows for an adequate representation of eukaryotic translation complexity and focuses on locations critical for translation regulation. We show how RDGs can be used for depicting translated regions, analysis of genetic variation and quantitative genome-wide data on translation for characterisation of regulatory modulators of translation.

## Motivation

Until relatively recently, available experimental evidence suggested that in eukaryotes each mRNA encoded only a single protein. Since only a single coding region was therefore expected to be translated this region was conventionally termed the CDS (CoDing Sequence). This view has been challenged by the development of the ribosome profiling technique which enables isolation and sequencing of RNA fragments protected by ribosomes and hence detection of regions being translated (1). Numerous ribosome profiling studies carried out in cells from a variety of eukaryotes unexpectedly revealed abundant translation outside of CDS regions. This included the translation of short sequences in the supposedly untranslated regions (UTRs) of mRNAs found upstream and downstream of the CDS, as well as in so-called non-coding RNAs, especially long non-coding RNAs (lncRNAs) (2–11). These studies also demonstrated the translation of N-terminally extended CDS regions due to initiation at upstream non-AUG start codons (12) or C-terminally extended due to stop codon readthrough (13). A certain group of eukaryotic organisms (ciliates *Euplotes*) were found to utilize ribosomal frameshifting in thousands of their genes (14). While most of these phenomena were first described prior to the advent of ribosome profiling (15–19), they were considered rare. Certainly, very few cases have been catalogued by reference gene annotation projects, and no conventional abstraction has been developed to represent this translation complexity in annotations, schematic scientific diagrams or analytical workflows. The lack of a formal framework for the representation of this complexity hampers our ability to generate accurate and biologically realistic annotations of translated sequences and to design mathematical models and computer simulations. In its absence, it is difficult or even impossible to quantitatively characterize multiple translation events and define their interrelationships.

To address this challenge, we developed a conceptually novel framework for abstract representation of translation complexity, which we term Ribosome Decision Graphs (RDGs). RDGs solve many problems, such as the representation of multiple translated regions in the same mRNAs and alternative decoding mechanisms producing multiple proteoforms. We show how RDGs can be used for the accurate depiction of productive and non-productive RNA translation (i.e. translation that does or does not lead to the production of a protein molecule), analysis of quantitative information on translation, and genetic variants affecting mRNA decoding.

### Open Reading Frames are inadequate to represent the complexity of mRNA translation

The development of a conventional abstraction is undermined by the ambiguity of the terms used to define translated regions. For example, while translated regions are often described as Open Reading Frames (ORFs) in literature or scientific discourse, gene annotation projects typically utilise only the term CDS, and only for regions considered to be protein-coding. Instead, an ORF would be regarded by implication as a potential translation that can be identified *in silico*. Here, we in effect consider three concepts in an attempt at unification: (1) that ORFs can be identified *in silico* whether or not they have evidence of translation, (2) that ORFs may undergo translation that does not lead to production of a stable, functional protein, and (3) that ORFs which are known to be translated into proteins should alone be considered as CDS. In other words, most CDS are ORFs, but not all ORFs are CDS. In general, there are two definitions of ORFs, start to stop (Start-Stop) and stop to stop (Stop-Stop)(20), as depicted in Figure 1A. Plotting the locations of potential start codons (usually AUGs) and stop codons in three reading frames is undoubtedly highly instrumental for examining potentially translated sequences. However, the common interpretation of nucleic acid sequences in terms of “translated ORFs” is superficial, frequently inaccurate, and often leads to confusion as illustrated in Figure 1B-D.

**Figure 1.**
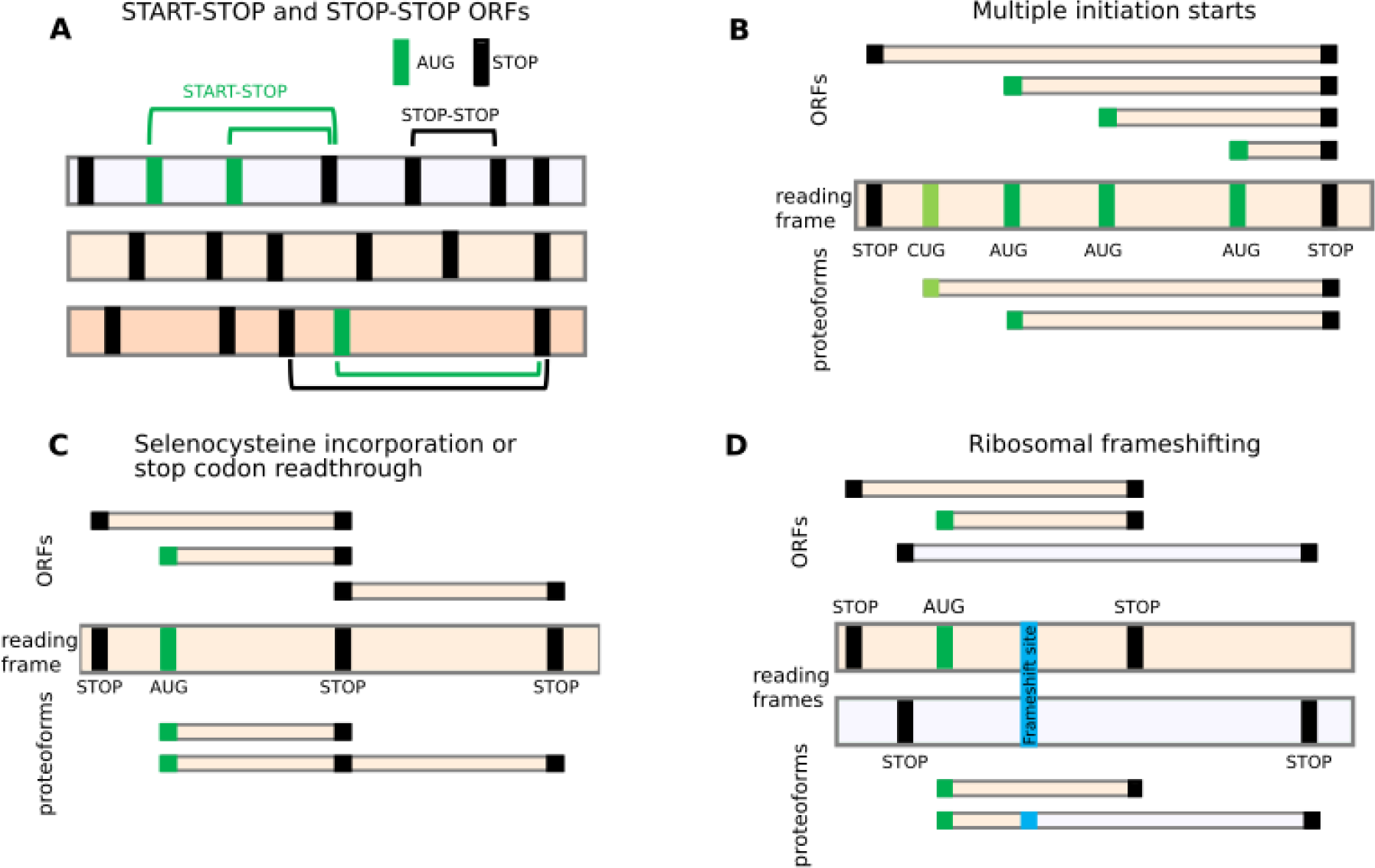
ORFs do not adequately represent translational complexity. **A**. Two formal definitions of ORFs. Three reading frames are shown as horizontal bars with vertical bars corresponding to AUG (green) or stop codons (black). Several examples of Start-Stop (green arcs) and Stop-Stop (black arcs) ORFs are shown. **B-D**. The relationship between ORFs (top) and expressed proteoforms (bottom) for mRNAs with different locations of starts and stops (middle). Only two relevant reading frames are shown for simplicity. **B**. An RNA encoding two proteoforms with alternative N-termini due to utilization of two start codons. Due to multiple potential AUG codons, there are many Start-Stop ORFs whose conceptual translation does not correspond to encoded proteoforms. Stop-Stop ORF does not reflect the existence of alternative proteoforms. **C**. In the case of stop codon readthrough or selenocysteine insertion, ribosomes may read through specific stop codons by incorporating an amino acid yielding a product (purple) that cannot be described as a product of a single ORF. **D**. Similarly, ribosomal frameshifting generates a trans-frame protein (blue) that does not represent a product of a single ORF.

Perhaps the most frequent source of alternative translation in many eukaryotes, including humans, is the multiplicity of translation initiation sites. It arises predominantly from two common mechanisms involved in the selection of translation initiation sites: leaky scanning and re-initiation. *Leaky scanning* refers to the inefficient recognition of a start codon by the ribosome resulting in the ribosome scanning complex scanning through the start codon and effectively ignoring it (21). Generally, ribosome scanning complexes assemble at the 5’ cap of RNAs and move along the transcript in the 3’ direction until they encounter a start site and initiate translation (22–24). However, recognition of start sites is a sequence-dependent stochastic process, in which usually only a proportion of scanning complexes finally initiate. Many factors play a role in determining the efficiency with which a ribosome initiates translation at a given codon. These include the identity of the codon and its surrounding sequence (known as the Kozak context) (25), as well as the dwell time of the scanning ribosome at that codon (26). Unless the combination of these factors is strictly optimal for initiation, at least a small fraction of scanning complexes will bypass the potential start site and continue scanning, allowing translation to be initiated further downstream. When a potential initiation site is a non-AUG codon or an AUG in a weak Kozak context, only a small proportion of scanning complexes will initiate translation. Thus, leaky scanning may result in the translation of different coding sequences using numerous initiation sites, while initiation at start codons in the same reading frame can give rise to proteoforms with alternative N-termini (PANTs) (Figure 1B). A potentially large number of start codons may be used to initiate translate within the same Stop-Stop ORF, as is the case with the well-explored human *PTEN* gene where functionally distinct extended proteoforms are produced from multiple non-AUG starts (27). Annotating all Start-Stop ORFs is problematic due to the large number of potential start codons, and in certain cases, such as Repeat Associated Non-AUG (RAN) translation (28,29), the exact position of the initiation site cannot even be easily identified.

In the case of stop codon readthrough (13,30) and selenocysteine incorporation (31,32) (Figure 1C), the CDS needs to be described as a fusion of a Start-Stop ORF with a Stop-Stop ORF (gene annotation projects currently resolve these cases by ‘re-writing’ the Stop codon or selenocysteine codon in the protein file, allowing them to code through). For programmed ribosomal frameshifting (Figure 1D), which is common in viruses but also infrequently occurs in cellular genes (33), the description of translation using ORFs would require the introduction of the location of the frameshift site as both start and stop codon. This could enable the designation of the trans-frame protein product as a fusion of the two such “ORFs”. In practice, gene annotation may instead introduce an artificial indel modification of the natural DNA/RNA sequences to yield a single contiguous ORF; for example, the [T] corresponding to human hg38 assembly chr19:2271440 nucleotide is deleted in both RefSeq (e.g. NM_004152.3) and Ensembl (e.g. ENST00000582888.8). To this end, existing gene annotation of in silico trans-frame translation may yield a protein sequence corresponding to the product generated in nature. However, it comes at the expense of producing an incorrect sequence of an mRNA molecule, which does not allow for the regulatory mechanism at play to be accurately represented.

The examples in Figure 1 are not exhaustive and there are other translation phenomena that cannot be easily described using ORFs, such as translational bypassing (34–36) and StopGo (also known as StopCarryOn or 2A) (37). Regardless of which ORF definition is used, the concept of a translated ORF is not adequate to represent the complexity of RNA translation.

### RNA translation is segmented

Ribosome profiling has revealed the existence of a large number of short translated sequences, currently termed small or short ORFs (smORFs, sORFs) or RiboSeq ORFs, as the term CDS is reserved for sequences encoding classical proteins (38). Many RiboSeq ORFs occur within the same RNA molecules. The lack of appropriate terminology reflecting the complexity of translation becomes even more evident when we consider the relationship between these translation segments. Upstream translation often influences downstream translation and this dependency is known to be utilised to regulate gene expression. For instance, many short translated regions upstream of CDSs (termed upstream ORFs, or uORFs) have been found to regulate translation by blocking ribosomes *via* sensing specific metabolites within the nascent peptide channel (39–43), reviewed in (44). This process is exemplified by translation regulation of the downstream CDS by a short uORF in vertebrate *AMD1* encoding Adenosyl-Methionin Decarboxylase, a key enzyme in polyamine biosynthesis. The uORF encodes a short peptide MAGDIS that stalls ribosomes through its interactions with the ribosome in the presence of polyamines (45). These stalled ribosomes prevent other ribosomes from binding and scanning downstream to initiate at *AMD1* CDS. Thus, the uORF provides a negative feedback control mechanism for *AMD1* expression, inhibiting its synthesis when polyamine concentration is high, but allowing for its synthesis when polyamine levels decrease (Figure 2A).

**Figure 2.**
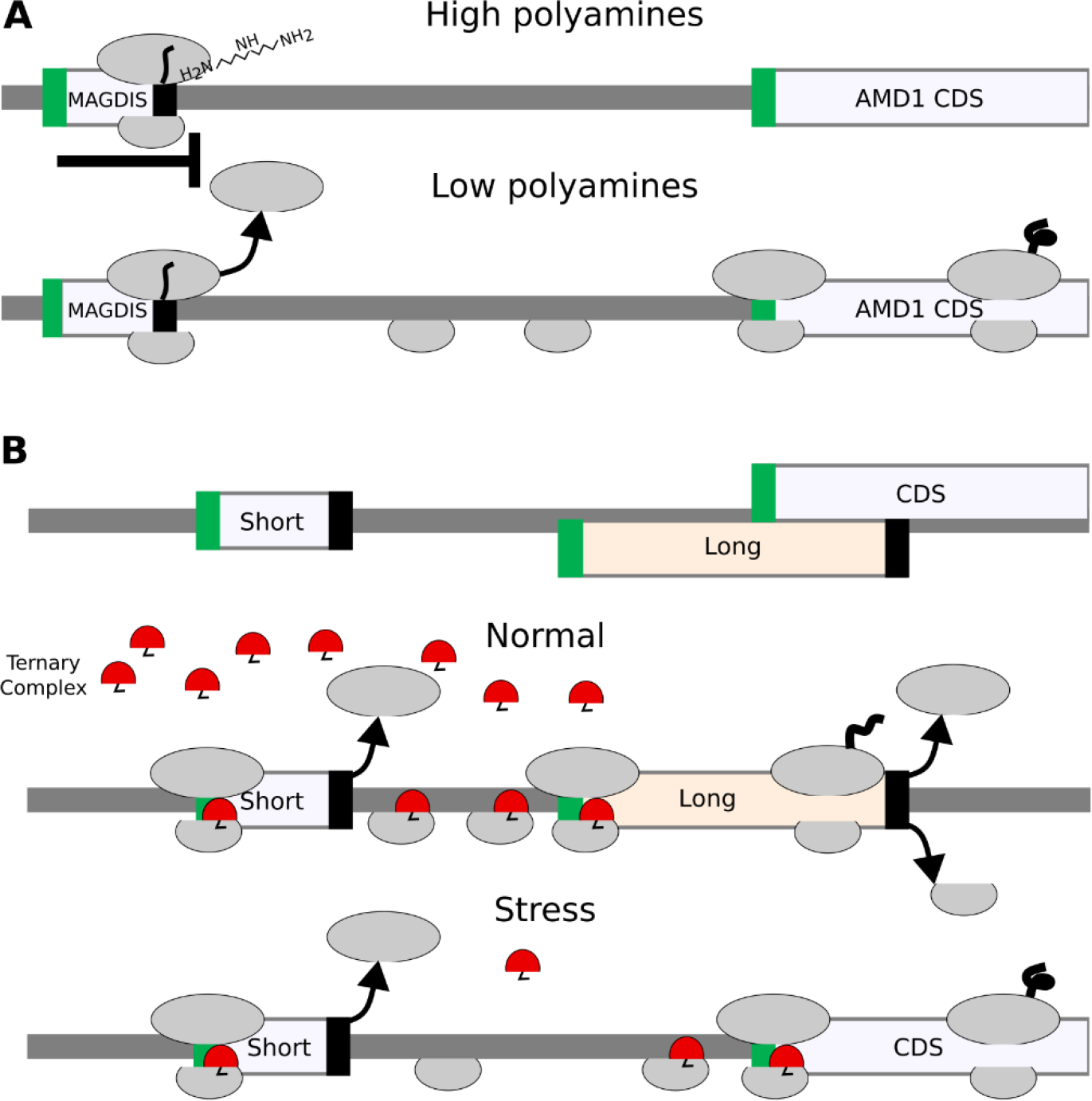
Relationship between translated segments within the same RNA. **A**. A schematic of metabolite-dependent translation regulation *via* ribosome arrest at uORF, such as in *AMD1* mRNA. In the presence of high concentration of polyamines ribosome with MAGDIS peptide stalls at the end of mRNA. **B**. A schematic of delayed re-initiation mechanism enabling translation of selected mRNAs during global translation suppression caused by eIF2alpha phosphorylation during Integrated Stress Response. When eIF2 is phosphorylated the concentration of the eIF2*tRNAi*GTP ternary complex decreases and it requires a longer time and distance for the scanning ribosome complex to acquire it. As a result the long uORF is bypassed and initiation occurs at CDS.

In addition to leaky scanning, re-initiation is another process impacting start codon selection. Translation re-initiation occurs when small ribosomal subunits remain bound to the mRNA after translation is complete and reinitiate downstream of the terminating stop codon. This is thought to be common after the translation of short ORFs as it takes time for initiation factors to dissociate from the ribosome. In this way, the ribosome may remain capable of initiation after translating a small number of codons, although other factors are known to contribute to this process, allowing for re-initiation in some instances even after the translation of long ORFs. The detailed molecular mechanisms of these processes are described in dedicated reviews (21,22,24,46–49). Re-initiation provides a platform for a rapid switch of gene expression on the translational level. Perhaps the most thoroughly studied is the case of delayed re-initiation (50–52) that protects translation of certain mRNAs (e.g. human A*TF4*, yeast *GCN4*) from downregulation during the Integrated Stress Response (ISR) (53,54). Under this condition, the reduced availability of the ternary complex (tRNAi* eIF2*GTP) increases the time required for post-terminating ribosomes to bind the ternary complex enabling re-initiation. Therefore, the level of stress determines the location of the start codon at which re-initiation occurs. Figure 2B provides a schematic illustrating this mechanism.

It Is unclear to what extent the translated products of such regulatory translation contribute to the functional cellular proteome beyond their potential contribution to the antigen pool, as many of them lack conservation at the protein level (38,55,56). Extreme cases of translation regulation without peptide synthesis are represented by minimal ORFs consisting of a start codon immediately followed by a stop codon. While they obviously do not produce any functional peptide, some of them do have regulatory potential as strong ribosome stalling sites (57).

It is clear that translation complexity requires a unified and comprehensive abstraction that would adequately represent all translated regions – not only those that encode classical proteins – and reflect their mechanistic interrelationships. Such representation should be convenient for use by both, human scientists when examining individual mRNA sequences and computer agents during programmatic analysis of large datasets.

### Ribosome Paths

The complex nature of translational events and regulatory processes reveals the need to consider the entire passage of an individual ribosomal complex containing the same small ribosomal subunit along the mRNA, from the moment of pre-initiation complex assembly at the 5’ cap (or IRES element) to the complete dissociation of both ribosomal subunits from the mRNA as a functional unit. We propose to term such a unit a Ribosome Path (RiboPath). It includes both regions that are scanned and those that are translated. As argued above, ORF is an inadequate descriptor of translated regions, and therefore we want to define and assign a new, unambiguous name to an entity denoting *translated region* as encompassing the entire sequence of RNA translated by a fully assembled elongating ribosome from initiation codon through termination and dissociation of the large ribosomal subunit. We term this region *translon*. It has already been suggested as a term specifying a unit of translation (58) but has not yet been adopted. The main advantage of *translon* over ORF is that it is not constrained by the sequence (specific codons as boundaries). It is based on the process of translation and therefore may incorporate a variety of decoding mechanisms such as ribosomal frameshifting, stop codon readthrough, translational bypassing, etc (59,60). The other term commonly used to indicate translated regions is *cistron*, e.g. polycistronic or monocistronic mRNAs. However, this term was originally defined genetically, different cistrons should be responsible for different phenotypes, and it is being used inconsistently in the literature.

To simplify the introduction of the RiboPath concept, for now, we only consider initiation and re-initiation as the mechanisms producing alternative proteoforms. We will exclude other translation mechanisms. Nevertheless, our framework can easily be extended to incorporate other translation mechanisms as we discuss later.

Figure 3 illustrates the RiboPath concept with an example of an mRNA encoding two proteoforms arising from alternative CUG and AUG initiation sites in one reading frame (cream) and a single upstream AUG codon in another frame (light lavender) as depicted in the ORF plot at the top. The corresponding translons are shown beneath. Alternative initiation and re-initiation allow the ribosome to pass through five different RiboPaths. The top RiboPath represents the ribosomes that initiate at the first AUG, but fail to reinitiate further downstream, resulting in a path with a single translon T1. The second path corresponds to the ribosome that successfully re-initiates downstream, thus containing two translons, T1 and T2. In the third RiboPath the ribosomes fail to initiate at the first AUG, but start translation at the CUG allowing for translon T3, which encodes an N-terminally extended proteoform relative to the product of translon T2. The fourth RiboPath corresponds to the ribosomes that fail to initiate at both the first AUG and the CUG but succeed at initiating at the second AUG so that its RiboPath consists of only one translon T2. Finally, the fifth RiboPath is unproductive and represents the ribosomes that have not initiated protein synthesis on this mRNA. The RiboPath presentation makes it clear that certain translons are mutually exclusive as they never occur on the same path, e.g. a single ribosome cannot translate T1 and T3.

**Figure 3.**
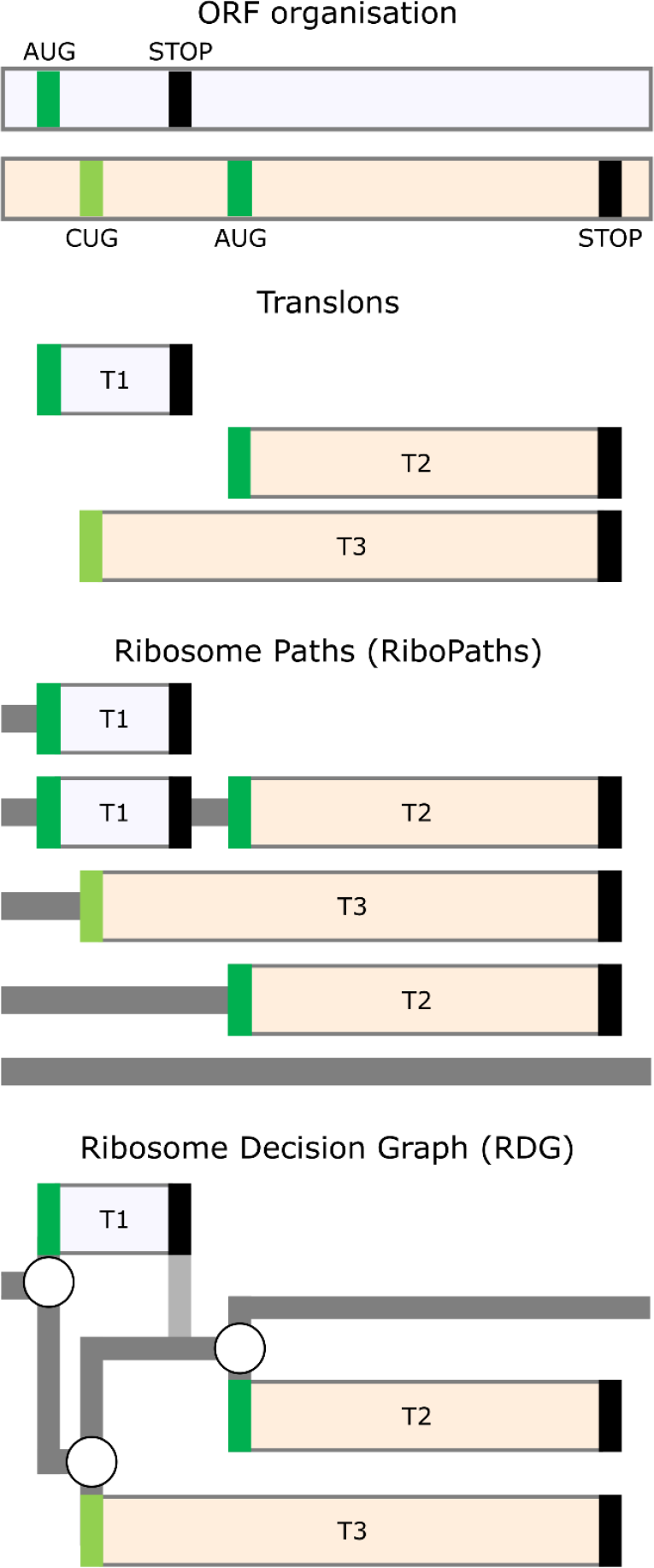
Ribosome paths through mRNA. **Top:** An example of an RNA with three start codons (green for AUG and light green for CUG) located in two different reading frames depicted as differentially shaded horizontal bars. The translation of this mRNA is represented as a set of translons **below**, Ribosome Paths **further below** and Ribosome Decision Graphs at the **bottom**. RNA regions scanned by the ribosome are shown in dark grey, vertical path in light grey represents the post-terminating small ribosome subunit that continues scanning and remains initiation-competent.

### Ribosome Decision Graphs

Once we represent the behaviour of translating ribosomes in terms of paths, it is only natural to further represent these in terms of graphs (Figure 3). The three initiation events in Figure 3 can be represented as branching points where the ribosome makes a “decision” of whether to initiate or not. We do not imply that ribosomes have free will; the decision is likely determined by the molecular composition and temporal thermodynamics of the local microstate. As in statistical mechanics, for practical purposes, it is appropriate to describe such decisions probabilistically, even if the underlying molecular processes are deterministic. The mRNA region engaged by ribosomes in Figure 3 can then be represented as a graph with three branching points. Stop codons in this graph are considered as deterministic ends of translons as we exclude the possibility of stop codon readthrough or re-initiation after long translons in our illustrative example. Following this notation, any translated RNA can be represented as a Ribosome Decision Graph (RDG).

As in the representation of translation using ORFs/CDS, RDGs may be either conceptual (representing potential) or real (e.g., experimentally supported). In the case of conceptual RDGs, all potential start codons in mRNAs could be used as branching points, e.g., all AUGs, all CUGs, etc. depending on the specific parameters of the model. Such conceptual RDGs would be very complex graphs with a large number of branching points and possible paths. They are not suitable for evaluation by humans, but provide a straightforward method for generating all theoretically possible products of RNA translation. This can be used for the subsequent mining of mass spectrometry datasets. A set of graphs with branching points sampled from the set of all possible branching points can be used to generate simulated ribosome profiling data. The comparison of simulated and real data would enable the determination of the best RDG fitting the experimental data, thus inferring the real branching points from the data. As exemplified further below, RDGs may also be useful for analysing the impact of genetic variation, because variants that change or introduce new branching points (start and stop codons, frameshifts, etc) would alter the RDG topology.

RDGs could also be used to annotate experimentally validated translations. In this case, only those translation events for which there is experimental evidence will be introduced as branching points. In most cases, these experimentally informed RDGs would be suitable for manual examination by researchers and would overcome the limitations of the data structures that are currently used for protein coding annotation.

### Implementations of RDGs

To illustrate how RDGs can be used to represent the impact of variation within 5’ leader sequences (a.k.a. 5’ UTR) on downstream translation we selected the *NF2* variant responsible for neurofibromatosis type 2 (61). The 5’ leader sequence of the *NF2* mRNA contains an AUG start codon followed by an in-frame AUG codon in a strong Kozak context. This suggests that few (if any) ribosomes reach the CDS start *via* leaky scanning. It is far more likely that CDS translation involves re-initiation at the CDS start as depicted in Figure 4A. A single base insertion variant was identified in two unrelated individuals in a cohort of 1,134 individuals diagnosed with neurofibromatosis type 2 (ENST00000338641:−66-65insT; GRCh37:chr22:29999922A >AT) (61). This insertion causes both a shift in the reading frame and the introduction of another AUG. The shift extends translons T1 and T2, abrogating the initiation of translon T3 corresponding to *NF2* CDS (Figure 4B).

**Figure 4:**
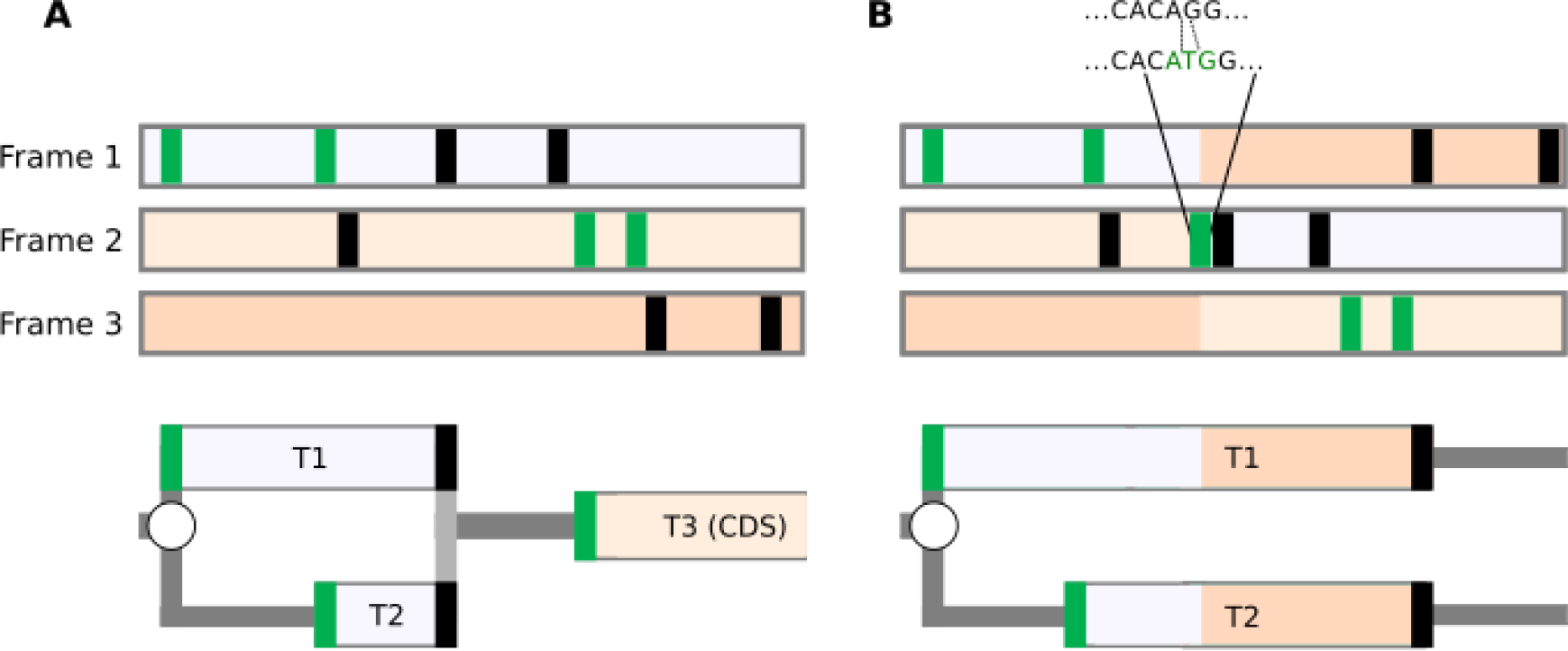
Representation of the effect of a genomic variant on CDS translation. Representations of the *NF2* mRNA for the reference sequence (**A**) and in the presence of the insertion variant (**B**). ORF organisation is shown at the **Top** with reading frames shaded differentially according to the reference sequence. AUG and stop codons are represented as green and black vertical bars, respectively. RDGs are shown at the bottom. Given the low probability of leaky scanning through the first two AUGs (see text), it is expected that the translon T3 corresponding to CDS cannot be translated in the pathogenic variant sequence.

To illustrate how RDGs can be used for the representation of real translation data we chose two simple examples, namely human *NRAS* and *NXT1* mRNAs. The criteria for this selection were the existence of only a single transcript per gene according to GENCODE v.42 (43), and the ribosome profiling supporting translation of only a single AUG-initiated translon in addition to the annotated CDS. Of note, translation of most human 5’ mRNA leaders is more complex, and therefore the advantages of using RDG representation for these are even greater, but may not be suitable for introducing this concept.

Examination of ribosome profiling data in Trips-Viz (62) (Figure 5A) for *NXT1* mRNA reveals translation of an upstream region in the −1 frame (blue translon) relative to the CDS (red translon). Similarly, examination of Trips-Viz data indicates translation upstream and in the +1 reading frame (red translon) relative to the annotated *NRAS* CDS (blue translon). For simplicity, the CDS starts are not depicted as a branching point and are considered to be 100% efficient translation initiation sites. As the translated regions in both graphs are overlapping, it is clear that the simultaneous translation of both translons by the same ribosome cannot occur, at least in the absence of 3’ to 5’ scanning of post-terminating ribosomes (63).

**Figure 5:**
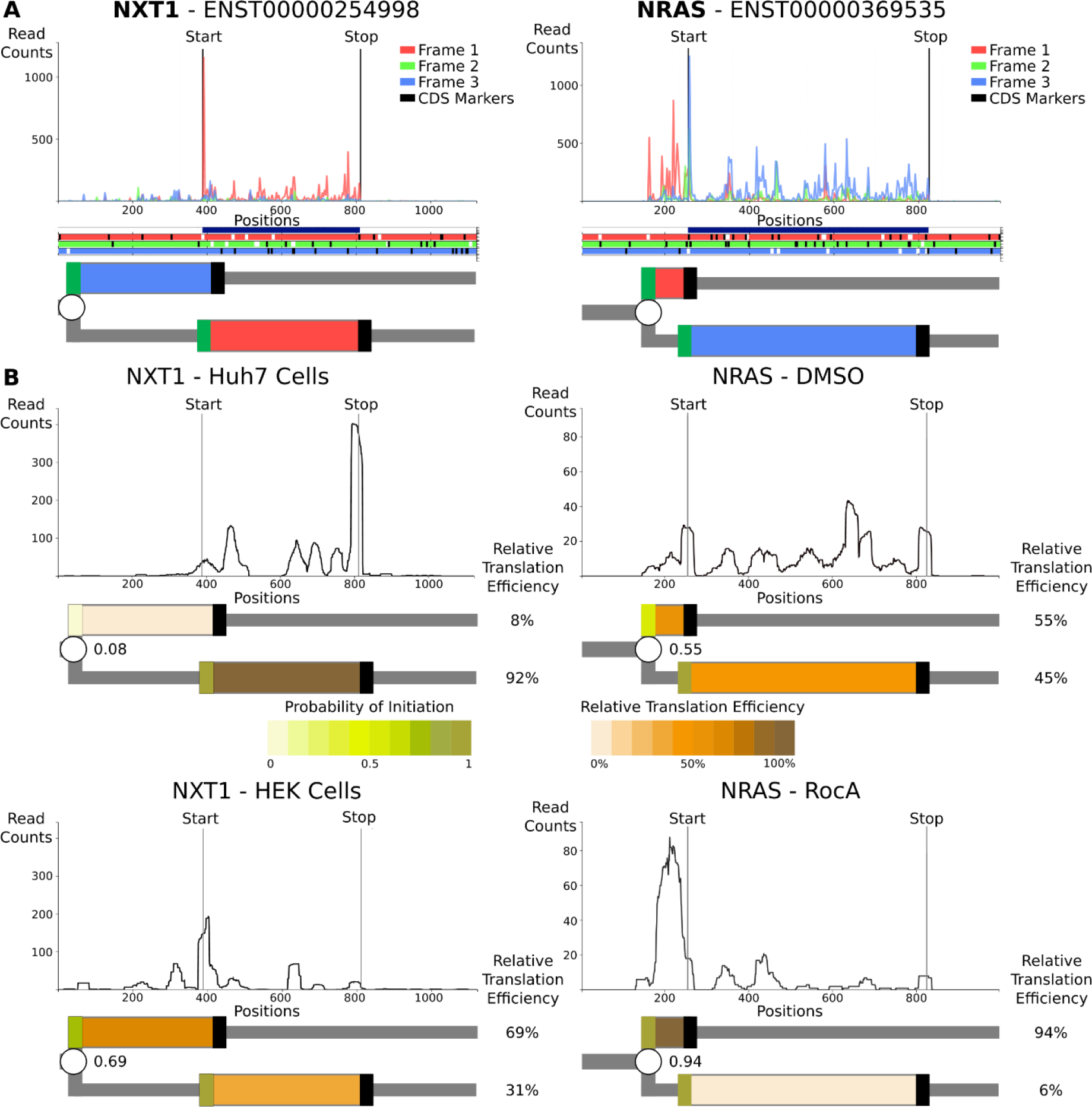
Representation of translation of human mRNAs. **A**. The top plots are A-site subcodon footprint densities for *NXT1* and *NRAS* mRNAs coloured to match the best-supported reading frame below in the ORF plots, in which AUGs are in white and stops are in black. Further below are RDG representations with translons coloured to match the translated frame. The CDS starts are treated deterministically and unproductive RiboPaths are not shown. **B**. Densities of ribosome A-site footprints obtained from ribosome profiling under different conditions (*NRAS*) or from different cells (*NXT1*). RDGs are shown below each density plot with translons coloured as a heatmap reflecting relative translation efficiencies. Branching points (starts) are also coloured as heatmaps reflecting the inferred probability of their initiation. The probabilities of translation initiation are shown as fractions decimals, while the relative translation efficiencies of each path are shown as percentages (%).

In addition to representing qualitative information, RDGs also enable a quantitative representation of translation regulation. Due to the leaky scanning mechanism of translation initiation, the efficiencies of CDS translation in these two examples directly depend on the efficiencies of the upstream starts, e.g., if all ribosomes initiated at the upstream starts, no CDS translation would be observed. The relative translation efficiencies of translons can be used to calculate the probabilities of initiation at the upstream starts (64). These probabilities may vary between different conditions or across different cell types due to a variety of mechanisms, such as global changes in the stringency of start recognition (65,66) or specific regulation of mRNA via ribosome sensing of particular metabolites through interaction with the nascent peptide (39–43). Using RDG representations in this way makes it easier to characterize the relationship between translation events that are regulated (*via* changes in probabilities at branching points) and the relative rates of translons product synthesis.

To illustrate this with real examples, we examined the translation of the above genes using different ribosome profiling datasets (Figure 5B). For *NRAS*, we used the dataset from cells treated with Rocaglamide A (RocA) and its untreated control (67). As can be seen in Figure 5B the silhouette of ribosome footprint density for the *NRAS* mRNA changes dramatically upon RocA treatment. These footprint densities can be used to calculate the relative translation efficiencies of *NRAS* translons and to derive the probability of translation initiation at the upstream starts (Methods). By showing the relative synthesis rates and initiation probabilities as heatmaps, the relationship between these two translons becomes apparent. RocA treatment greatly increases translation initiation probability at the upstream start, most likely *via* the ability of RocA to clamp eIF4A to mRNAs containing specific sequence motifs (67), which then reduces the downstream CDS translon. In the case of *NXT1*, we examined data obtained in two different cell lines, HeLa (68) and Huh7 (69). The silhouettes of ribosome footprint densities for *NXT1* mRNA are markedly different, as can be seen in Figure 5B. The RDG visualization of these differences in ribosome footprint densities pinpoints the upstream start as the pivotal element of cell-specific regulation of *NXT1* translation. For HeLa samples, the translation initiation at the upstream start is highly efficient, making the upstream start predominant. In contrast, for Huh7 samples efficiency at this start is much lower and, consequently, the CDS translon is predominant. The reasons for these cell-specific differences are beyond the scope of this work, but several mechanisms may be responsible, including different levels of translation factors that recognize translation initiation starts (70).

### Future prospects

One of the attractive features of the RDG concept is its expandability. In the RDG examples above, we limited branching points only to starts where initiation and re-initiation events can occur. The most basic information for generating RDGs requires only locations of starts in a transcript because in-frame stop codons are identifiable from the sequence and are treated deterministically as the ends of translons. Therefore, the example shown in Figure 1B can be represented as a short array:

#### starts(x_1_;x_2_;x_3_;x_4_)

while the example in Figure 3 as

#### starts(x_1_;x_2_;x_3_)

However, the concept can be extended to incorporate annotations for any non-deterministic translation events. For example, to capture such phenomena as stop codon readthrough or selenocysteine insertion (Figures 1C and 6A), an addition of a new type of branching point would be necessary: a stop codon, at which the ribosome could either terminate or incorporate an amino acid and continue translation.

**Figure 6:**
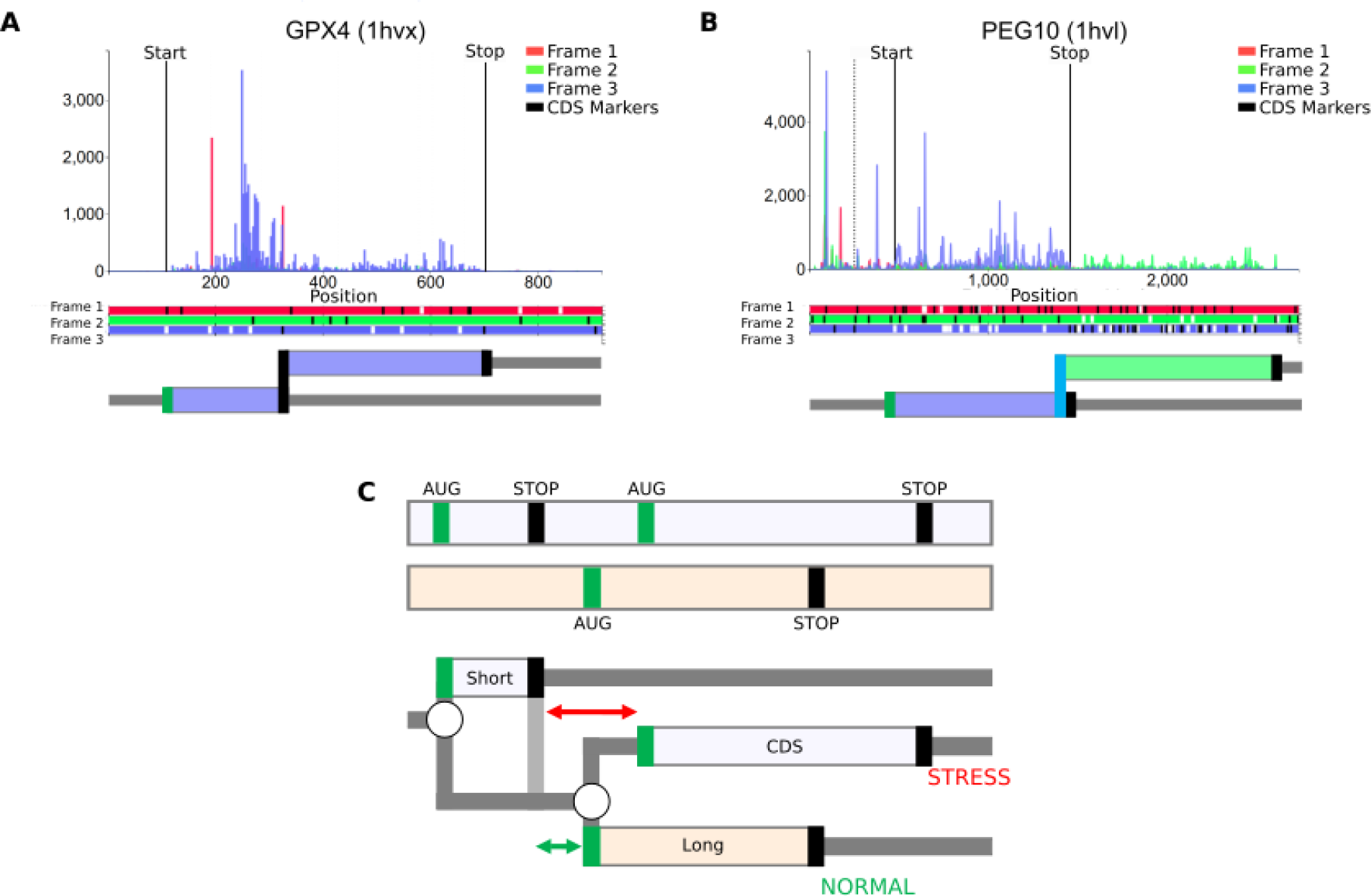
RDG representations of special cases. (**A**) mRNA encoding selenoprotein GPX4 and (**B**) PEG10 mRNA requiring ribosomal frameshifting for its expression. Trips-Viz screenshots of ribosomal profiling density for these mRNAs are at the top with ORF plots beneath and RDG representations further below. **C. Top**: An ORF plot containing three translons (protein-coding CDS and two uORFs, short and long), which is a minimal requirement for the mechanism of delayed re-initiation. **Below**: The corresponding RDG with green and red representing the predominant ribosome paths for normal and stress conditions, respectively. Arrows indicate the distance sufficient for scanning ribosomes to bind ternary complex under normal (green) or stress (red) conditions.

#### starts(x_1_); secs(y_1_)

A frameshifting site would be yet another type of branching point where the ribosome could either continue translation in the same frame or shift to one of the alternative reading frames. For example, the case of ribosomal frameshifting (Figures 1D and 6B) can be represented as

#### starts(x_1_); shifts(i_1_,s_1_)

where shifts need to be assigned two coordinates, the coordinate of the last codon in the initial frame ***i***_***1***_ and the coordinate of the first codon in the shifted frame ***s***. Accordingly, ***s***_***1***_ – ***i***_***1***_ = 1 would correspond to +1 frameshifting, ***s***_***1***_ – ***y***_***1***_ = −1 to −1 frameshifting, and so on. This notation is compatible not only with common types of frameshifting but also with extreme events such as bypassing in bacteriophage T4 gene *60* (34), where 50 nucleotides are skipped by the ribosome without translation, i.e. expressed as ***shifts(i***,***s)*** where ***s*** – ***i*** = 50.

Algorithms interpreting such annotations will be backward and forward-compatible since older algorithms could simply ignore new types of branching points.

Despite the apparent simplicity of RDGs notations, it would be naive to expect that it can represent a full range of translational mechanisms. For example, in the case of a delayed re-initiation mechanism (50–52) that makes translation of certain mRNAs resistant to global downregulation during Integrated Stress Response (ISR), it is not sufficient to simply add a stop codon as a branching point, allowing either ribosome dissociation from mRNA or re-initiation downstream. This is because the reduced availability of the ternary complex (tRNAi*eIF2*GTP) increases the time required for the post-terminating ribosomes to bind the ternary complex, thereby enabling re-initiation (Figure 2B). Thus, it is not the probability of re-initiation, but the location of the start at which re-initiation will occur that changes during ISR. However, even in this case, the RDG concept can be useful to illustrate the mechanism, as shown in Figure 6C. It is also conceivable to extend the concepts of RDGs with parameters linking scanning distance to re-initiation probability.

An important shortcoming of the presented solution is the difficulty of its application to genomic loci encoding multiple transcript isoforms. The purpose of RDGs is to represent molecular events that take place during the translation of a single mRNA molecule. Therefore, a single RDG can only be applied to a single mRNA sequence. However, the concept of representing biological sequences as graphs is gaining momentum with splice graphs for representing alternative splicing (71) and variation graphs for representing pangenomes (72). Therefore, we envision that the RDG concept will fit into the emerging bioinformatics infrastructure of hierarchical representation of biological sequences as graphs, from genome to transcriptome to translatome.

## Methods

We used GENCODE v42 (73) as a source of transcriptome annotation when searching for suitable loci and GENCODE v25 when producing visualizations in Trips-Viz (62). Pre-processed alignments from datasets GSE70211, GSE79664, and GSE94454 obtained in relevant studies (67–69) were obtained from Trips-Viz. The translation efficiencies of examined regions were calculated as the number of ribosome footprints uniquely aligning to individual translons normalised over the length of translons used for the mapping. Five first codons of translons were excluded from mapping to avoid the distortions introduced by the high peaks at the starts. The probability of translation initiation was then calculated as *p=R*_*u*_*/(R*_*u*_*+R*_*d*_), where *R*_*u*_ and *R*_*d*_ are translation efficiencies of the upstream and downstream translons.

## Acknowledgements

J.A.S.T. is supported by Science Foundation Ireland Centre for Research Training in Genomics Data Science [18/CRT/6214], M.S. and J.K are supported by Poland National Science Centre [UMO-2021/41/B/NZ2/03036] to J.K. N.W. is supported by a Sir Henry Dale Fellowship jointly funded by the Wellcome Trust and the Royal Society (220134/Z/20/Z) and research grant funding from the Rosetrees Trust (PGL19-2/10025), H.T. is supported by Irish Science Foundation Frontiers for the Future award ([20/FFP-A/8929] to P.V.B. J.M.M. is supported by the Wellcome Trust (grant number 108749/Z/15/Z), the National Human Genome Research Institute (NHGRI) of the US National Institutes of Health (NIH) under award number 2U41HG007234 and the European Molecular Biology Laboratory (EMBL). E.V. is funded by the Research Council of Norway (#314216). P.V.B. also wishes to acknowledge Investigator in Science Award by the SFI-HRB-Wellcome Trust Biomedical Research Partnership ([210692/Z/18/]. The content is solely the responsibility of the authors and does not necessarily represent the official views of the National Institutes of Health. Ensembl is a registered trademark of EMBL.

